# Protective effects of low-intensity pulsed ultrasound on mandibular condylar cartilage exposed to mechanical overloading

**DOI:** 10.1101/325886

**Authors:** Mutsumi Fujita, Minami Sato-Shigeta, Hiroki Mori, Akihiko Iwasa, Nobuhiko Kawai, Ali H Hassan, Eiji Tanaka

## Abstract

The aim of this study was to examine the role of low-intensity pulsed ultrasound (LIPUS) exposure in the onset and early progression of temporomandibular joint (TMJ) osteoarthritis (TMJ-OA) induced by mechanical overloading. Fifteen-week-old male Wistar rats were divided into two experimental groups and a control group (n = 5 each). In the experimental groups, both TMJs were subjected to mechanical overloading by forced mouth opening using a jaw-opening device for 3 h/day for 5 continuous days. After mechanical overloading, TMJs in one experimental group were exposed to LIPUS for 20 min/day. After the experiments, mandibles were resected from the rats, and the condyles were processed. The bones were analyzed using high-resolution microcomputed tomography (micro-CT). The resected TMJs were also subjected to histological analysis and immunohistochemical staining. Micro-CT images of the mandibular condyle showed severe subchondral trabecular bone loss in the experimental group with overloading. Treatment with LIPUS after overloading resulted in decreased subchondral trabecular bone resorption. In TMJ sections from the experimental group with overloading, cell-free regions and proteoglycan loss characterized the cartilage degradation; LIPUS exposure restricted these changes in the mandibular condyle. Furthermore, the number of tartrate-resistant acid phosphatase-positive osteoclasts in the mineralized layer of the condylar cartilage increased after mechanical overloading and decreased after LIPUS treatment. Our findings suggest that LIPUS exposure after mechanical TMJ overloading downregulates subchondral trabecular bone resorption and proteoglycan loss in the mandibular condylar cartilage. Thus, it may prove to be protective effects of LIPUS exposure on onset and early progression of TMJ-OA induced by mechanical overloading.

## Introduction

Synovial joints play a key role in relatively large bone movements induced by surrounding muscle forces [1]. The bone ends come together within a fibrous joint capsule. The synovium, a metabolically active tissue, covers the inner lining of this joint capsule. The bone ends are covered with a thin layer of dense connective tissue known as the articular cartilage [2]. Ligaments, tendons, and other soft tissues around the joint cavity provide stability to the joint and maintain appropriate alignment of the articulating bone ends during motion [2]. Daily activity accompanies joint motion, resulting in joint loads. The temporomandibular joint (TMJ), a diarthrodial synovial joint, enables large relative movements between the temporal bone and the mandibular condyle [3, 4]. The integrity of the joint is maintained intrinsically by a fibrous capsule with ligamentous thickenings and extrinsically by accessory ligaments, both of which restrict movement at the extremes of the mandibular range of motion and, consequently, have limited influence on the mechanics of normal symmetrical function [5, 6]. Within the joint, the articular surfaces of the condyle and articular eminence are covered by a thin fibrocartilaginous layer with a very low coefficient of friction [5].

The most common pathology affecting TMJ is degenerative joint disease, also known as osteoarthritis (OA). The hallmarks of OA include collapse of the articular cartilage, osteophyte formation, and subsequent joint space narrowing [7]. TMJ-OA is characterized by degradation of the mandibular condylar cartilage due to mechanical overloading [8–10]. Mechanical overloading of the mandibular condylar cartilage increases the expression of interleukin-1β (IL-1β) [11], an inflammatory cytokine closely associated with the progression of TMJ-OA [10, 12]. IL-1β markedly stimulated osteoclastic bone resorption by enhanced both osteoclast formation and function [13]. Because the fibrocartilage covering both the TMJ condyles and glenoid fossa is avascular, its cells have limited ability for self-repair [14, 15], similar to hyaline cartilage in other synovial joints [16]. Therefore, once cartilage breakdown starts, TMJ-OA can be crippling, leading to various morphological and functional deformities. This suggests the importance of suppressing cartilage degradation during the early stages of this disease. Although moderate physical exercise has been associated with delayed onset and slow progression of OA in humans [17, 18], no treatment remedy has been developed for severe OA.

Low-intensity pulsed ultrasound (LIPUS) is an acoustic radiation source with an intensity of less than 100 mW/cm^2^. It has been widely accepted as a therapeutic, operative, and diagnostic tool in the medical field. Previous in vivo studies have demonstrated that LIPUS can promote bone healing, remodeling, and regeneration and augment osteogenesis at the distraction site. It is generally accepted that LIPUS has no deleterious or carcinogenic effects. Moreover, LIPUS exposure has no thermal effects inducing biological changes in living tissues. Previously, we reported that LIPUS reduces inflammation and promotes regeneration in various conditions involving injured soft tissues, such as synovitis of the knee joint, injury of the skeletal muscle, and sialadenitis [19–22]. Therefore, we hypothesized that LIPUS may be a good treatment strategy for inflammatory diseases affecting the articular cartilage, such as OA. The aim of the present study was to examine the role of LIPUS exposure in the onset and early progression of TMJ-OA induced by mechanical overloading.

## Materials and methods

### Animals

Fifteen-week-old male Wistar rats were used in this investigation. They were kept in plastic cages, in which the wall and floor were smooth, and fed a stock diet with water ad libitum. The animals were randomly divided into one control group and two experimental groups of 5 animals each. In the two experimental groups, both TMJs were subjected to mechanical overloading for 5 continuous days. TMJs in the OL group received only mechanical overloading for 5 days, while those in the OL+LIPUS group were exposed to LIPUS for 20 min/day after mechanical overloading. In the control group, TMJs were not subjected to any form of mechanical overloading, although the same anesthesia schedule was maintained. All procedures performed in this study were approved by the Tokushima University Animal Care and Use Committee (Permit No: T27-93).

### Application of Mechanical Stress

In the experimental groups, both TMJs were subjected to mechanical overloading by forced mouth opening using a jaw-opening device for 3 h/day. The device maintained a mouth opening of 20 mm. During the mouth-open period, the rats were anesthetized with an intra-abdominal injection of sodium pentobarbital (Nembutal; Dinabott, Osaka, Japan) at a dose of 50 mg/kg body weight.

### LIPUS

In the OL+LIPUS group, LIPUS was applied with a modified version of the clinical device, Osteotron V (ITO Co., Tokyo, Japan) after mechanical overloading. The ultrasound exposure system was equipped with a circular surface transducer with a cross-sectional area of 5.0 cm^2^. The ultrasound head exhibited a mean beam nonuniformity value of 3.6 and an effective radiating area of 4.1 cm^2^. An ultrasound signal was transmitted at a frequency of 1.5 MHz, with a spatial average intensity of 30 mW/cm^2^ and a pulse rate of 1:4 (2 ms on and 8 ms off). Both TMJs were exposed to LIPUS for 20 min/day during the 5-day experimental period.

### Microcomputed Tomography (Micro-CT)

After the experiments, the rats were sacrificed with an overdose of anesthesia. The animals in the control group were sacrificed after 5 days. The mandibles were resected, and the condyles were carefully separated from the surrounding soft tissues and fixed in 70% ethanol overnight. The bones were then analyzed using high-resolution micro-CT (SkyScan 1176 scanner, Buruker, Billeriea, MA) and its analysis software. Briefly, images were acquired at 50 kV and 500 μA. During scanning, the samples were tightly covered with plastic wrap to prevent movement and dehydration. Thresholding was applied to the images to segment the bone from the background. The resolution of the micro-CT images was 9 μm/pixel. The ratio of the bone volume (BV) to the trabecular volume (TV), trabecular thickness (Tb.Th), and trabecular number (Tb.N) were calculated.

### Tissue Preparation and Histological Analysis

After the animals were sacrificed, both TMJs were resected, fixed in 10% buffered paraformaldehyde, and decalcified with 10% EDTA at 4°C for 8 weeks. Thereafter, they were embedded in paraffin, and serial sections (7 μm) were cut in the sagittal plane. The sections were stained with hematoxylin and eosin (HE) for histological examination and stained with 0.1% Safranin-O and 0.02% fast green for the detection of cartilage and proteins, respectively. In addition, toluidine blue staining was used to detect proteoglycans in the condylar cartilage.

A modified Mankin scoring system [23] was used to assess the degree of cartilage degeneration. The Safranin-O-stained sections were used for scoring the features of cartilage disease, including changes in cellularity, structural abnormalities, and safranin-O uptake as a measure of glycosaminoglycan distribution and loss. The sections were blindly examined by three independent experts.

### Tartrate-Resistant Acid Phosphatase (TRAP) Staining

TRAP activity was measured to identify the characteristics of osteoclast lineage cells according to the method of Minkin [24]. The staining medium consisted of naphtol AS-MX phosphate (Sigma Chemical Co., St. Louis, MO, USA) as a substrate, Fast Red Violet LB Salt (Sigma Chemical Co., St. Louis, MO, USA) as a coupler, and 50-mM sodium tartrate (Wako, Osaka, Japan). Hematoxylin was used for counterstaining. Negative staining was performed without substrate.

### Immunohistochemistory

To investigate the expression of aggrecan, type II collagen (Col2a1), matrix metalloproteinase (MMP)-9, and MMP13 in the condylar cartilage, immunohistochemical staining was performed using various primary antibodies (Immuno Biological Laboratories, Fujioka, Japan). After the sections were deparaffinized and blocked in, they were incubated overnight at 4°C in a humid atmosphere with various primary antibodies diluted in phosphate-buffered saline /0.1% bovine serum albumin. Immunostaining was performed using a Histofine simple stain kit (Nichirei, Tokyo, Japan). Briefly, after blocking endogenous peroxidase activity with 0.3% hydrogen peroxide in methanol, nonspecific binding of the antibody was blocked by incubating the section for 30 min with Non-Specific Staining Blocking Reagent (Dako, Carpinteria, Calif., USA). After washing, the sections were incubated with the corresponding secondary antibodies for 1 h at room temperature, followed by mounted. The sections were examined under a BioRevoBZ-9000 microscope (KEYENCE, Osaka). Negative controls were stained with nonimmune immunoglobulin G.

### Statistical Analysis

All values are expressed as means and standard deviations. Significant differences in experimental data were analyzed using one-way analysis of variance, followed by the Turkey-Kramer test as a post hoc test to examine mean differences at a significance level of 5%.

### Results

The health status of the experimental groups was similar to those in the control group through the experimental period.

### Micro-CT

Micro-CT images showed severe subchondral trabecular bone loss in the OL group, but not in the control group (Fig 1). In contrast, treatment with LIPUS after overloading reduced the amount of subchondral trabecular bone resorption in the OL+LIPUS group. Furthermore, BV/TV, Tb.Th, and Tb.N in different areas of the condylar subchondral bone were significantly lower in the OL group than in the control group (P < 0.01), whereas BV/TV and Tb.Th were significantly higher in the OL+LIPUS group than in the OL group (P < 0.05). Although Th.N was also higher in the OL+LIPUS group than in the OL group, the difference was not significant. Although all values were higher in the control group than in the OL+LIPUS group, the differences were not significant.

**Fig 1.**
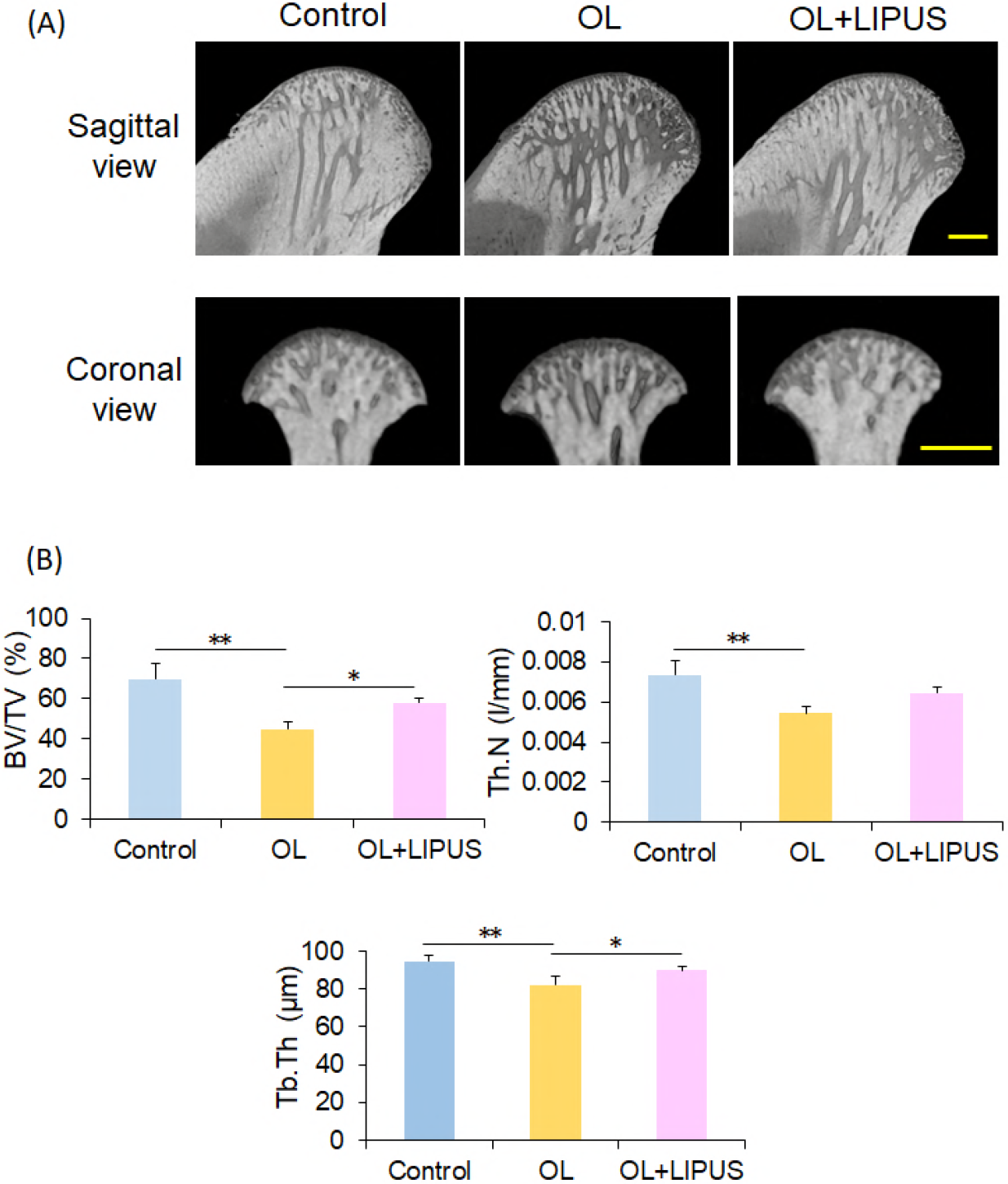
Effects of low-intensity pulsed ultrasound (LIPUS) exposure on experimental temporomandibular joint arthritis in rats. (A) Three-dimensional reconstruction of mandibular condyles from the control and experimental mice, with representative sagittal and coronal sections obtained from microcomputed tomography scans of the condyles. (B) The trabecular bone volume (BV) was calculated using a representative sagittal section and expressed as the BV/trabecular volume (TV) ratio. * P < 0.05; ** P < 0.01 as per Tukey-Kramer tests (n = 5 for each group); Scale bars = 100 μm in for the sagittal view and 1000 μm in for the coronal view.

### Histology

HE staining of the mandibular condyles from the control group showed that the surface of the condylar cartilage was smooth, with obviously distinguished cell layers (Fig 2). In contrast, the mandibular condyles from the OL group showed marked OA-like lesions, including a decrease in the thickness of the cartilage layer, irregularities in chondrocyte alignment, compressive necrosis of chondrocytes, and hyalinization of the cartilage matrix in the proliferating, mature, and hypertrophic cell layers. However, the amount of damage was markedly lesser in the condylar cartilages from the OL+LIPUS group.

**Fig 2.**
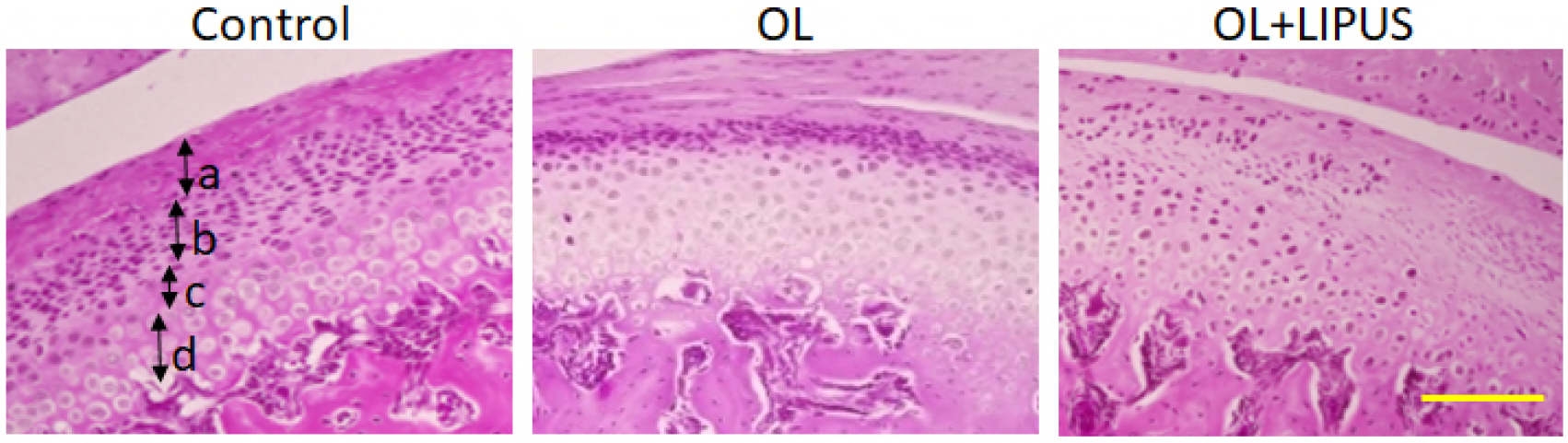
Hematoxylin and eosin (HE)-stained sections of the mandibular condylar cartilage from experimental [temporomandibular joint arthritis induced by mechanical overloading, with or without low-intensity pulsed ultrasound (LIPUS) exposure] and control rats. a: fibrous cell layer, b: proliferating cell layer, c: mature cell layer, d: hypertrophic cell layer; Scale bar = 100 μm.

Safranin-O-stained sections showed that proteoglycans were abundant in the deep layer of the mandibular condylar cartilages from the control group (Fig 3A). However, the amount of proteoglycans was significantly decreased in the samples from the OL group. The modified Mankin scores confirmed that overloading caused significant (P < 0.01) changes in structural characteristics that paralleled the progression of OA (Fig 3C). LIPUS exposure after overloading significantly (P < 0.01) lowered the modified Mankin score relative to that for the OL group, although the scores were still significantly lower in the control group than in the OL+LIPUS group (P < 0.01). Attenuated toluidine blue staining was observed in the condylar cartilages from the OL group; this reduction in staining was restricted after LIPUS exposure (Fig 3B). The significantly reduced amount of proteoglycans in the condylar cartilage (P < 0.05) after overloading was restored to the level observed for the control group after LIPUS exposure (Fig 3D).

**Fig 3.**
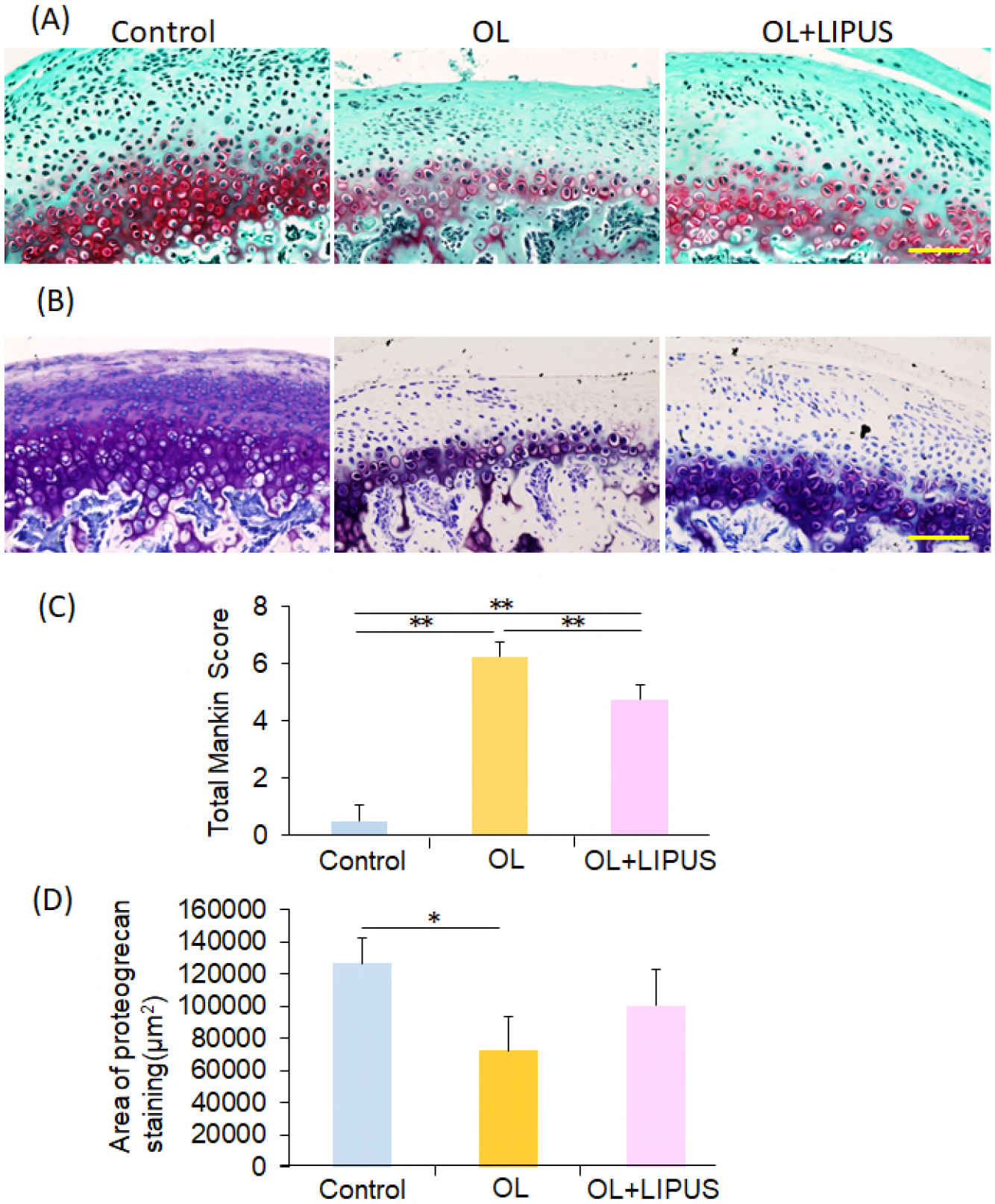
Histochemically stained sections of the mandibular condylar cartilage from experimental [temporomandibular joint arthritis induced by mechanical overloading, with or without low-intensity pulsed ultrasound (LIPUS) exposure] and control rats. (A) Safranin-O and fast green staining. (B) Toluidine blue staining. (C) Comparison of the findings of proteoglycan staining among the three groups. (D) Histological grading according to modified Mankin scores for the mandibular condylar cartilages obtained from the three groups. Data are expressed as means and standard deviations. * P < 0.05; ** P < 0.01 as per Tukey-Kramer tests (n = 5 for each group); Scale bars = 100 μm.

### Number of TRAP-positive osteoclasts

In the control group, the number of TRAP-positive osteoclasts in the mineralized layer of the condylar cartilage was 5.6 ± 0.9 (Fig 4A, B). This number was 13.2 ± 3.0 after mechanical overloading and 10.0 ± 1.6 after LIPUS exposure. Thus, the number of TRAP-positive osteoclasts was significantly smaller in the control group than in the OL (P < 0.01) and OL+LIPUS (P < 0.05) groups.

**Fig 4.**
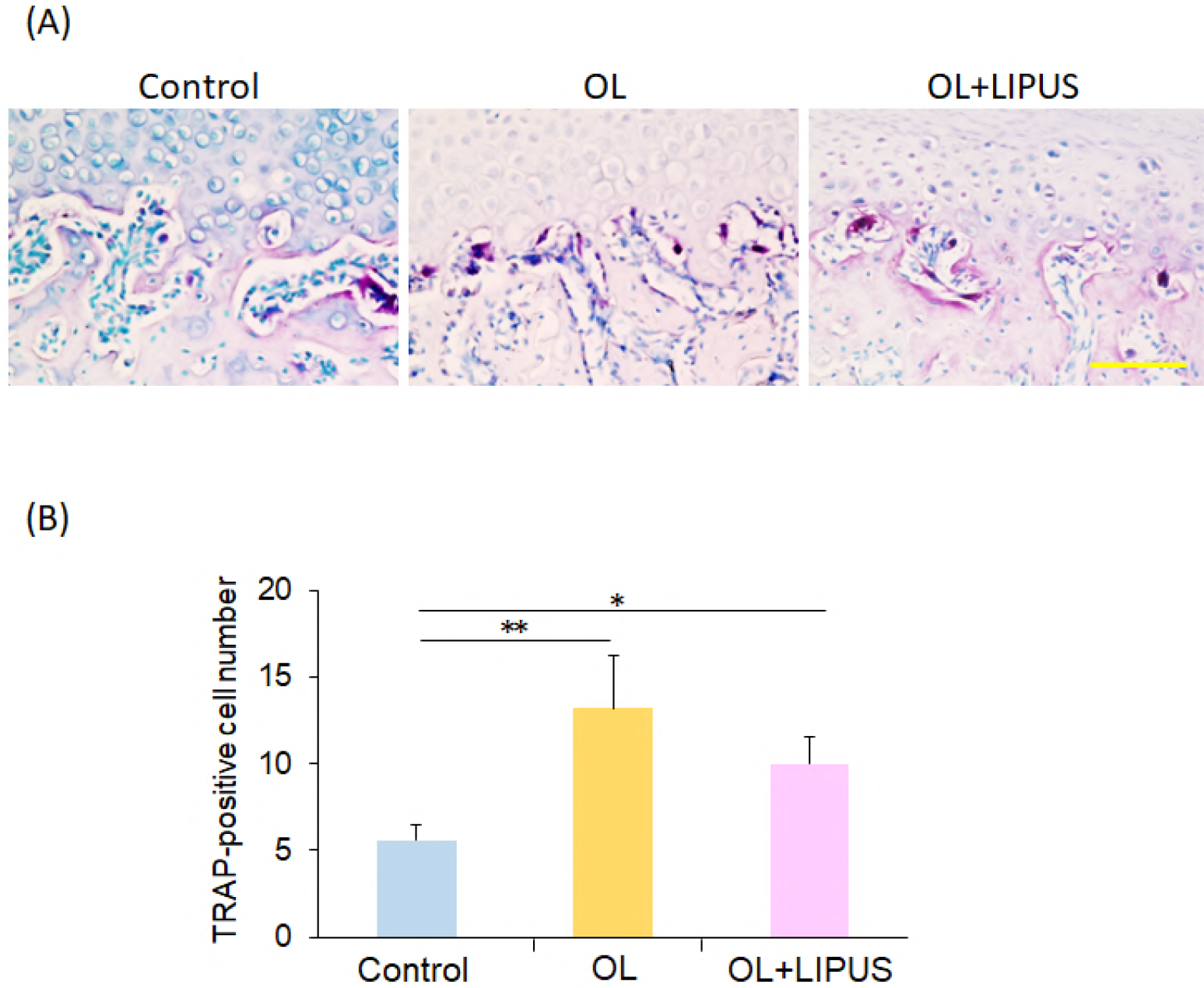
Tartrate-resistant acid phosphatase (TRAP) staining for the subchondral bone adjacent to the mandibular condylar cartilage in experimental [temporomandibular joint arthritis induced by mechanical overloading, with or without low-intensity pulsed ultrasound (LIPUS) exposure] and control rats. (A) TRAP staining. (B) Comparison of the number of TRAP-positive cells among the three groups. * P < 0.05; ** P < 0.01 as per Tukey-Kramer tests (n = 5 for each group); Scale bar = 100 μm.

### Immunohistochemical analysis

Immunohistochemistry showed more MMP9- and MMP13-positive cells and lesser Col2a1- and aggrecan-positive cells in the OL group than in the control group (Fig 5A-D). LIPUS exposure downregulated the number of MMP9- and MMP13-positive cells and upregulated the number of Col2a1- and aggrecan-positive cells in the condylar cartilages from the OL+LIPUS group. These data show that LIPUS exposure downregulates the expression of the cartilage destruction factor.

**Fig 5.**
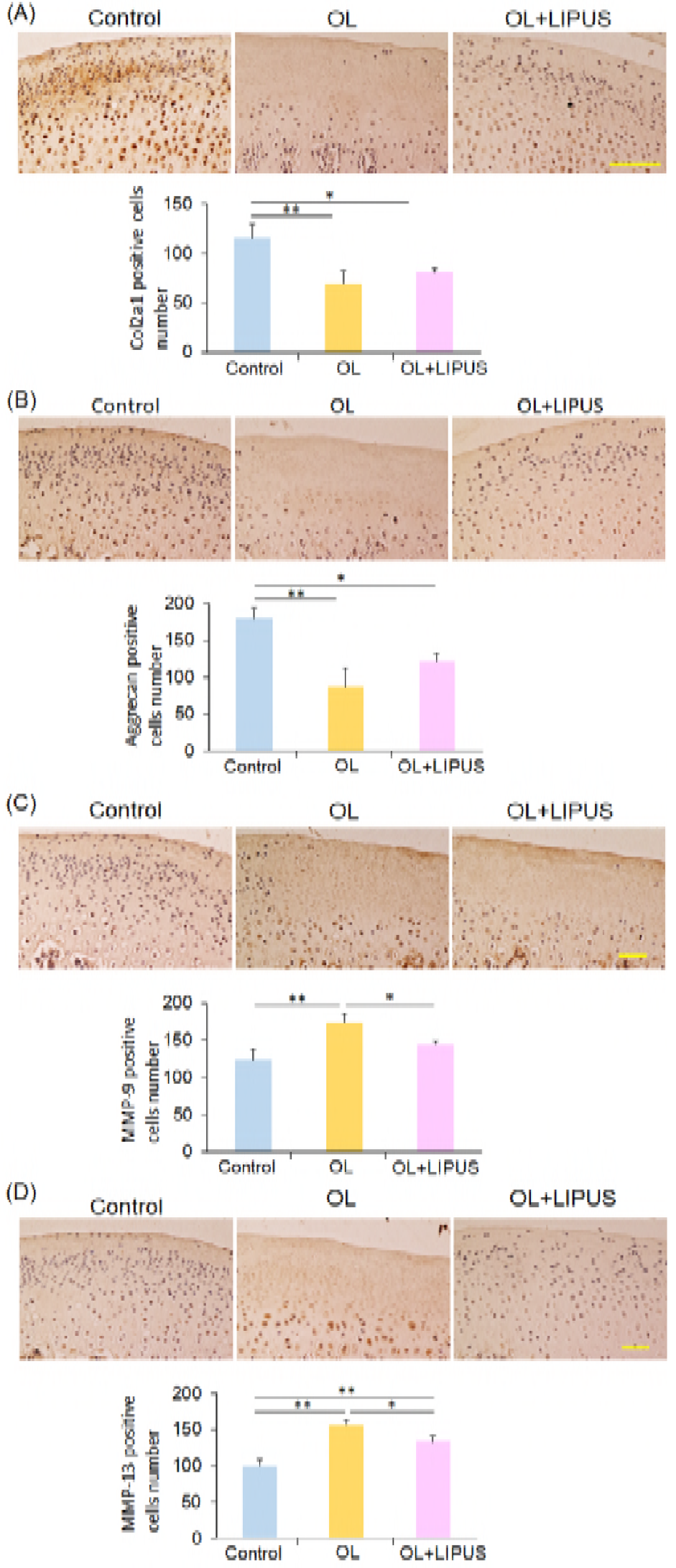
Immunohistochemical staining for aggrecan, type II collagen, and matrix metalloproteinase (MMP)-9 and MMP13 in mandibular condylar cartilage samples from experimental [temporomandibular joint arthritis induced by mechanical overloading, with or without low-intensity pulsed ultrasound (LIPUS) exposure] and control rats. (A) Aggrecan. (B) Collagen type II (Col2a1). (C) MMP9. (D) MMP13. * P < 0.05; ** P < 0.01 as per Tukey-Kramer tests (n = 5 for each group); Scale bars = 100 μm.

The number of Col2a1- and aggrecan-positive cells was significantly larger in the control group than in the OL (P < 0.01) and OL+LIPUS (P < 0.05) groups (Fig 5A, B). The OL group showed significantly more MMP9- and MMP13-positive cells than did the control (P < 0.01) and OL+LIPUS groups (P < 0.05).

## Discussion

In the present study, we examined the role of LIPUS exposure in the onset and early progression of TMJ-OA induced by mechanical overloading and found that LIPUS exposure after mechanical TMJ overloading downregulates subchondral trabecular bone resorption and proteoglycan loss in the mandibular condylar cartilage, resulting in protective effects of LIPUS on the mandibular condylar bone resorption and cartilage degradation.

OA, the most common joint disease, is characterized by joint pain and stiffness with subsequent disability. The prevalence rates for symptomatic knee OA in individuals aged 60 years or more range from 10% to 13% [25]. OA is also the most common joint pathology affecting TMJ; among patients with TMJ disorders, 11% showed symptoms of TMJ-OA [26]. Management strategies for TMJ-OA may be divided into noninvasive and invasive strategies, and for end-stage disease, salvage modalities must be considered [9]. Although conventional treatment modalities for OA are pharmacological as well as nonpharmacological (regulation of daily activities, education, and exercise), no remedy has been able to revive cartilage degeneration. Therefore, ultrasound therapy aimed at tissue repair and regeneration has garnered considerable attention.

Previously, several studies showed the efficacy of LIPUS exposure to pathological chondrocytes derived from diseased knee cartilage. Uddin et al. [27] indicated the potential of LIPUS therapy to prevent cartilage destruction through mechanical stimulation, thereby inhibiting the catabolic action of IL-1β and stimulating chondrocyte migration, proliferation, and differentiation. Nishida et al. [28] also stated that CCN protein 2 (CCN2) production with chondrocyte differentiation was regulated by mitogen-activated protein kinase (MAPK) pathways activated by LIPUS-induced Ca^2+^ influx. Furthermore, we investigated the effects of LIPUS exposure on IL-1β-induced COX-2 expression in cultured chondrocytes derived from porcine mandibular condyles and emphasized that LIPUS exposure inhibited IL-1β-induced COX-2 expression via the integrin β1 receptor, followed by the phosphorylation of extracellular signal-related kinase (ERK) 1/2 [29]. However, the effects of LIPUS on the mandibular condylar cartilage in patients with TMJ-OA induced by mechanical overloading *in vivo* remain unclear. To the best of our knowledge, this is the first study investigating the effects of LIPUS on the onset and early progression of TMJ-OA induced by mechanical overloading.

In the present study, exposure of overloaded condylar cartilages to LIPUS attenuated cartilage degradation, reduced the number of osteoclastic cells, and reduced the expression of MMP9 and MMP13. In addition, the amount of proteoglycans in condylar cartilages treated with LIPUS after forced mouth opening was restored to the same level observed in the control group. These results imply the inhibitory effects of LIPUS on osteoclastic bone resorption and condylar cartilage degradation in patients with TMJ-OA induced by mechanical overloading. Jia et al. [30] evaluated the efficacy of focused LIPUS therapy as a noninvasive modality for knee OA in a double-blind placebo-controlled trial including 106 patients with bilateral knee OA. The authors found that LIPUS is a safe and effective modality for pain relief and improvement of the knee joint function in OA patients. Similarly, the present study indicates that LIPUS is a noninvasive treatment modality that prevents mandibular condylar resorption and cartilage degradation at the onset and during the early progression of TMJ-OA.

Because of its limited capacity for regeneration, articular cartilage requires structural and metabolic support after traumatic and/or chronic damages. However, available techniques for cartilage regeneration have either been invasive or ineffective. Recently, Yilmaz et al. [31] conducted a randomized controlled trial of LIPUS treatment in a rat model of knee OA and demonstrated that extracorporeal therapy and LIPUS exposure have systemic proliferative and regenerative effects on cartilage and tissue. Our results revealed that treatment with LIPUS upregulated the expression of aggrecan and type II collagen, which was reduced after mechanical overloading. However, the levels were still significantly lower than those in the control group. Micro-CT showed that BV/TV and Tb.Th were significantly greater for mandibular condyles treated with LIPUS after mechanical overloading than for overloaded condyles without LIPUS treatment; moreover, these parameters were almost the same as those for untreated condyles. Furthermore, the modified Mankin score for the mandibular condyles from the OL+LIPUS group was significantly lower than that for the condyles from the OL group, although the score for the condyles from the OL+LIPUS group was significantly higher than that for the condyles from the control group. These results indicate that LIPUS contributes, to some extent, to condylar cartilage repair and regeneration in patients with TMJ-OA induced by mechanical overloading. Further studies are needed to identify the maximal effects of LIPUS on cartilage regeneration.

Cartilage is a highly mechanoresponsive tissue. Chondrocytes function as mechanosensors and undergo a series of complex changes, including proliferation and metabolic alteration, as targets of external biomechanical and biochemical stimuli [27]. Therefore, mechanical stimulation such as LIPUS exposure may inhibit the catabolic action of IL-1β and stimulate chondrocyte migration, proliferation, and differentiation. IL-1β is a known inflammatory mediator that acts through the nuclear factor (NF)-kB pathway to induce the expression of several genes that are upregulated in cartilage affected by OA, such as *IL-1β, IL-6, IL-8,* and *MMP13* [32]. Several reports suggest that LIPUS exposure activates integrins on the cell membrane that act as mechanoreceptors to promote the attachment of various focal adhesion adaptor proteins [33]. We also demonstrated that LIPUS exposure caused significant upregulation of phosphorylated focal adhesion kinase (FAK) in knee synovial joint cells, and that the inhibition of FAK phosphorylation led to significant downregulation of MAPK phosphorylation [34, 35]. Nishida et al. [28] also reported that gene expression of chondrocyte differentiation markers and connective tissue growth factor (CTGF) production were increased in cultured chondrocytes treated with LIPUS, and that Ca^2+^ influx and phosphorylation of p38 MAPK and ERK1/2 were increased by LIPUS treatment. Moreover, exposure of cementoblasts to LIPUS mediated cell metabolism through the MAPK pathway because LIPUS enhanced the protein expression of ERK1/2; moreover, evidence suggested that treatment with an MEK1/2 inhibitor suppresses the upregulation of *Cox-2* mRNA expression induced by LIPUS [36]. Because the MAPK pathway is considered a general pathway involved in cell proliferation [37], the biological effects of LIPUS exposure on chondrocytes may be particularly promoted by the integrin/FAK/MAPK pathways. Further studies should evaluate the detailed mechanisms underlying the effects of LIPUS on articular cartilage affected by TMJ-OA.

## Conclusions

The results of this study suggest that LIPUS treatment attenuates cartilage degeneration, decreases the number of osteoclastic cells, and protects the onset and early progression of TMJ-OA induced by mechanical overloading. Moreover, it restores the expression of aggrecan and type II collagen after an initial decrease induced by mechanical overloading. Therefore, LIPUS exposure may represent an effective treatment strategy for TMJ-OA induced by mechanical overloading. These findings may have important implications for future ultrasound research, particularly in terms of the management of TMJ-OA.

## Acknowledgments

We are grateful to Tsukasa Kurahashi, Nobuyasu Yamanaka, and Atsushi Chuma for providing the ultrasound devices and technical assistance during the experiments.

## Author Contributions

MF, MSS, ET: Conception and design MF, MSS, AI: Conduct of experiments

Analysis and interpretation of data: MF, MSS, MH, AI

Senior advisors: MSS, NK, ET

Drafting of the Article: MF, AHH, ET.

## References

1. Widegren U, Wretman C, Lionikas A, Hedin G, Henriksson J. Influence of exercise intensity on ERK/MAP kinase signalling in human skeletal muscle. Pflugers Archiv - Eur J Physiol. 2000; 441(2-3):317–22.

2. Warwick R, Williams PL. Gray’s Anatomy. Philadelphia, PA: Saunders Co., 1973.

3. Rees LA. The structure and function of the mandibular joint. Br Dent J. 1954;96:125–33.

4. Scapino RP, Obrez A, Greising D. Organization and function of the collagen fiber system in the human temporomandibular joint disk and its attachments. Cells Tissues Organs. 2006;182(3-4):201–25.

5. Tanaka E, Kawai N, Tanaka M, Todoh M, van Eijden T, Hanaoka K, et al. The frictional coefficient of the temporomandibular joint and its dependency on the magnitude and duration of joint loading. J Dent Res 2004;83(5):404–7.

6. Kawai N, Tanaka E, Takata T, Miyauchi M, Tanaka M, Todoh M, et al. Influence of additive hyaluronic acid on the lubricating ability in the temporomandibular joint. J Biomed Mater Res. 2004;70A(1):149–53.

7. Arends RH, Karsdal MA, Verburg KM, West CR, Bay-Jensen AC, Keller DS. Identification of serological biomarker profiles associated with total joint replacement in osteoarthritis patients. Osteoarthritis Cart. 2017;25(6);866–77.

8. Leonardi R, Lo Muzio L, Bernasconi G, Caltabiano C, Piacentini C, Caltabiano M. Expression of vascular endothelial growth factor in human dysfunctional temporomandibular joint discs. Arch Oral Biol. 2003;48(3):185–92.

9. Tanaka E, Detamore MS, Mercuri LG. Degenerative disorders of the temporomandibular joint: Etiology, diagnosis, and treatment. J Dent Res. 2008;87(4):296–307.

10. Kuroda S, Tanimoto K, Izawa T, Fujihara S, Koolstra JH, Tanaka E. Biomechanical and biochemical characteristics of the mandibular condylar cartilage. Osteoarthritis Cart. 2009;17(11):1408–15. doi: 10.1016/j.joca.2009.04.025.

11. Yoshida H, Fujita S, Nishida M, Iizuka T. Immunohistochemical distribution of lymph capillaries and blood capillaries in the synovial membrane in cases of internal derangement of the temporomandibular joint. J Oral Pathol Med. 1997;26(8):356–61.

12. Ghassemi-Nejad S, Kobezda T, Rauch TA, Matesz C, Glant TT, Mikecz K. Osteoarthritis-like damage of cartilage in the temporomandibular joints in mice with autoimmune inflammatory arthritis. Osteoarthritis Cart. 2011;19(4):458–65. doi: 10.1016/j.joca.2011.01.012.

13. Kusano K, Miyaura C, Inada M, Tamura T, Ito A, Nagase H, et al. Regulation of matrix metalloproteinases (MMP-2, -3, -9, and -13) by interleukin-a and interleukin-6 in mouse calvaria: association of MMP induction with bone resorption. Endocrinology. 1998;139(3):1338–45.

14. Scheven BA, Man J, Millard JL, Cooper PR, Lea SC, Walmsley AD, et al. VEGF and odontoblast-like cells: stimulation by low frequency ultrasound. Arch Oral Biol 2009;54(2):185–91.

15. Neumann E, Lefèvre S, Zimmermann B, Gay S, Müller-Ladner U. Rheumatoid arthritis progression mediated by activated synovial fibroblasts. Trends Mol Med. 2010;16(10):458–68. doi: 10.1016/j.molmed.2010.07.004.

16. Iwamoto M, Ohta Y, Larmour C, Enomoto-Iwamoto M. Toward regeneration of articular cartilage. Birth Defects Res C Embryo Today. 2013;99(3):192–202. doi: 10.1002/bdrc.21042.

17. Toyoda T, Seedhom BB, Kirkham J, Bonass WA. Upregulation of aggrecan and type II collagen mRNA expression in bovine chondrocytes by the application of hydrostatic pressure. Biorheology. 2003;40(1-3):79–85.

18. Griffin TM, Guilak F. The role of mechanical loading in the onset and progression of osteoarthritis. Exerc Sport Sci Rev. 2005;33(4):195–200.

19. Nagata K, Nakamura T, Fujihara S, Tanaka E. Ultrasound modulates the inflammatory response and promotes muscle regeneration in injured muscles. Ann Biomed Eng. 2013;41(6):1095–105. doi: 10.1007/s10439-013-0757-y.

20. Nakamura T, Fujihara S, Katsura T, Yamamoto K, Inubushi T, Tanimoto K, et al. Effects of low-intensity pulsed ultrasound on the expression and activity of hyaluronan synthase and hyaluronidase in IL-1β-stimulated synovial cells. Ann Biomed Eng. 2010;38(11):3363–70. doi: 10.1007/s10439-010-0104-5.

21. Nakamura T, Fujihara S, Yamamoto-Nagata K, Katsura T, Inubushi T, Tanaka E. Low-intensity pulsed ultrasound reduces the inflammatory activity of synovitis. Ann Biomed Eng. 2011;39(12):2964–71. doi: 10.1007/s10439-011-0408-0.

22. Sato M, Kuroda S, Mansjur KQ, Ganzorig K, Nagata K, Horiuchi S, et al. Low-intensity pulsed ultrasound rescues insufficient salivary secretion in autoimmune sialadenitis. Arthritis Res Ther. 2015;17:278. doi: 10.1186/s13075-015-0798-8.

23. Xu L, Flahiff CM, Waldman BA, Wu D, Olsen BR, Setton LA, Li Y. Osteoarthritis-like changes and decreased mechanical function of articular cartilage in the joints of mice with the chondrodysplasia gene (cho). Arthritis and Rheumatism 2003;48:2509e2518.

24. Minkin C. Bone acid phosphatase: tartrate-resistant acid phosphatase as a marker of osteoclast function. Calcif Tissue Int. 1982;34(3):285–90.

25. Zhang Y, Jordan JM. Epidemiology of osteoarthritis. Clin Geriatr Med. 2010;26(3):355–69. doi: 10.1016/j.cger.2010.03.001.

26. Mejersjö C, Hollender L. TMJ pain and dysfunction: relation between clinical and radiographic findings in the short and long-term. Scand J Dent Res. 1984;92(3):241–8.

27. Uddin SM, Richbourgh B, Ding Y, Hettinghouse A, Komatsu DE, Qin YX, et al. Chondro-protective effects of low intensity pulsed ultrasound. Osteoarthritis Cart. 2016;24(11):1989–98. doi: 10.1016/j.joca.2016.06.014.

28. Nishida T, Kubota S, Aoyama E, Yamanaka N, Lyons KM, Takigawa M. Low-intensity pulsed ultrasound (LIPUS) treatment of cultured chondrocytes stimulates production of CCN family protein 2 (CCN2), a protein involved in the regeneration of articular cartilage: mechanism underlying this stimulation. Osteoarthritis Cart. 2017;25(5):759–69. doi: 10.1016/j.joca.2016.10.003.

29. Iwabuchi Y, Tanimoto K, Tanne Y, Inubushi T, Kamiya T, Huang YC, et al. Effects of low-intensity pulsed ultrasound on the expression of cyclooxygenase-2 in mandibular condylar chondrocytes. J Oral Facial Pain Headache. 2014;28(3):261–8. doi: 10.11607/ofph.1156.

30. Jia L, Wang Y, Chen J, Chen W. Efficacy of focused low-intensity pulsed ultrasound therapy for the management of knee osteoarthritis: a randomized, double blind placebo-controlled trial. Sci Rep. 2016;6:35453. doi: 10.1038/srep35453.

31. Yilmaz V, Karadaş Ö, Dandinoğlu T, Umay E, Çakçı A, Tan AK. Efficacy of extracorporeal therapy and low-intensity pulsed ultrasound in a rat knee osteoarthritis model: A randomized controlled trial. Eur J Rheumatol. 2017;4(2):104–8. doi: 10.5152/eurjrheum.2017.160089.

32. Goldring MB. Chondrogenesis, chondrocyte differentiation, and articular cartilage metabolism in health and osteoarthritis. Ther Adv Musculoskelet Dis. 2012;4(4):269–85. doi: 10.1177/1759720X12448454.

33. Lal H, Verma SK, Smith M, Guleria RS, Lu G, Foster DM, et al. Stretch-induced MAP kinase activation in cardiac myocytes: differential regulation through β1-integrin and focal adhesion kinase. J Mol Cell Cardiol. 2007;43(2):137–47.

34. Sato M, Nagata K, Kuroda S, Horiuchi S, Mansjur K, Nakamura T, et al. Low-intensity pulsed ultrasound activates integrin-mediated mechanotransduction pathway in synovial cells. Ann Biomed Eng. 2014;42(10):2156–63. doi: 10.1007/s10439-014-1081-x.

35. Tanaka E, Kuroda S, Horiuchi S, Tabata A, El-Bialy T. Low-intensity pulsed ultrasound in dentofacial tissue engineering. Ann Biomed Eng. 2015;43(4):871–86. doi: 10.1007/s10439- 015-1274-y.

36. Rego EB, Inubushi T, Kawazoe A, Tanimoto K, Miyauchi M, Tanaka E, et al. Ultrasound stimulation induces PGE_2_ synthesis promoting cementoblastic differentiation through EP2/EP4 receptor pathway. Ultrasound Med Biol. 2010;36(6):907–15. doi: 10.1016/j.ultrasmedbio.2010.03.008.

37. Cowan KJ, Storey KB. Mitogen-activated protein kinases: new signaling pathways functioning in cellular responses to environmental stress. J Exp Biol. 2003;206(7):1107–15.

